## Supplementary Figures for "Identifying Space-Resolved Proteins of the Murine Thymus, by Combining MALDI Mass Spectrometry Imaging and Proteomics"

#### Supplementary Methods

##### PepBridge.py program description

PepBridge.py is a Python program designed to map MALDI-identified m/z values with LCMS m/z values. The inspiration for this program came from our efforts to match MALDI with LCMS m/z values using the Excel's MATCH function. This function is typically used to determine the position of a lookup value within a column table, returning its relative position. However, the MATCH function in Excel has a limitation – it only returns the position of the first exact match encountered.

PepBridge.py overcomes this limitation by seamlessly associating MALDI-identified m/z values with their corresponding LCMS values within a user-defined threshold.

At its core, PepBridge.py reads two CSV files containing MALDI and LCMS m/z values and generates a consolidated file containing all LCMS rows matching the MALDI m/z value.

The program's logic is straightforward: it begins by loading MALDI and LCMS data files in CSV format, extracting the respective masses. Initially stored as Python lists, these masses are later converted to NumPy arrays for enhanced search efficiency. After extracting the masses from both LCMS and MALDI data files, the program iterates over the MALDI masses, identifying matches in the LCMS data using the get_hits method. This method effectively retrieves all LCMS matching masses within a specified threshold. To ensure a well-organized structure, the program adopts and object-oriented approach. PepBridge.py comprises of three main classes: MatchingMZ.py, MALDI_LCMS.py and Config_file.py.

Config_file.py primarily serves as a class for reading and manipulating configuration files in INI format. It offers functionality and methods for reading INI files, allowing users to specify various parameters such as the location of LCMS or MALDI output files, weather m/z values should be rounded to the nearest integer before matching, and which columns should be read from the files (such as m/z, average, control and experimental values). This flexibility is invaluable, particularly when dealing with CSV containing numerous columns. Furthermore, the [ordered_columns] field within the INI file enables users to specify the desired order in which columns should be printed, adding an extra layer of customization to the output.

MALDI_LCMS is the class that triggers the reading of INI configuration files and initializing the corresponding LCMS and MALDI objects. It load the MALDI/LCMS CSV data into memory and provides functionality to order the file columns according to the user's specified order, as well as to extract the columns holding the MALDI m/z columns (rounded or not) used for a precise or approximate matching among the LCMS masses (m/z).

The MatchingMZ class serves as the workhorse, performing the LCMS and MALDI class initialization and providing functionality to perform the m/z mapping between MALDI to LCMS data and store the matches into a file.

### Supplementary Material

### Figures

### Figure S1

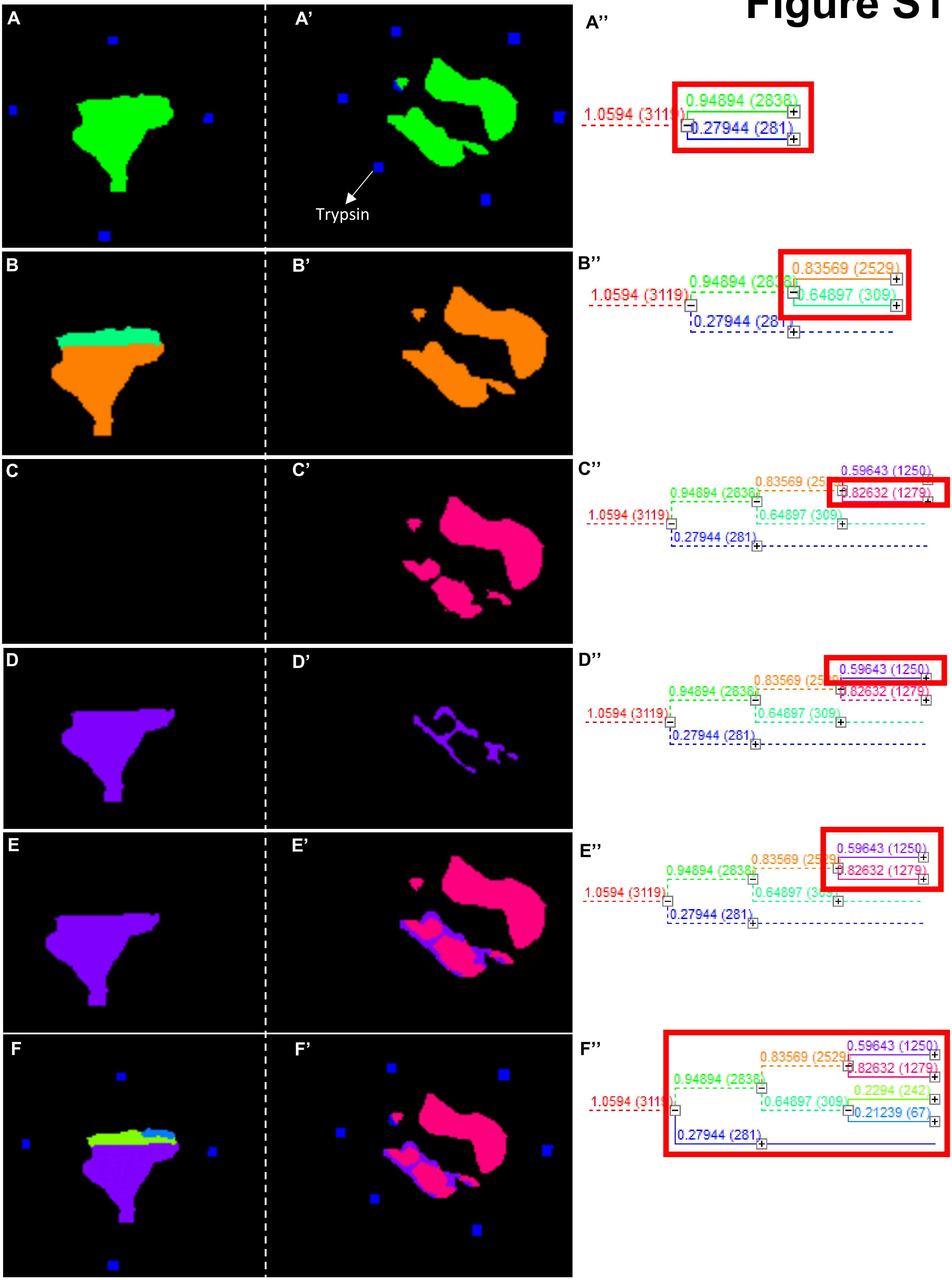


**Supplementary Figure S1.** Unsupervised hierarchical clustering of similar spectra represented as nodes in the dendogram. Each node is assigned a specific color that generates a pixelated image of the same color to the location of the node of clustered spectra on the tissue. For example, the blue and red nodes boxed in red in **(A")** correspond to images shown on the left in (**A** and **A'**). The blue node contains 281 spectra that correspond to trypsin while the green node contains 2838 spectra that correspond to the tissue. Nodes from **B"to G"** were selected to show that the hierarchical clustering works for compartmentalization of the thymus tissue.

### Figure S2

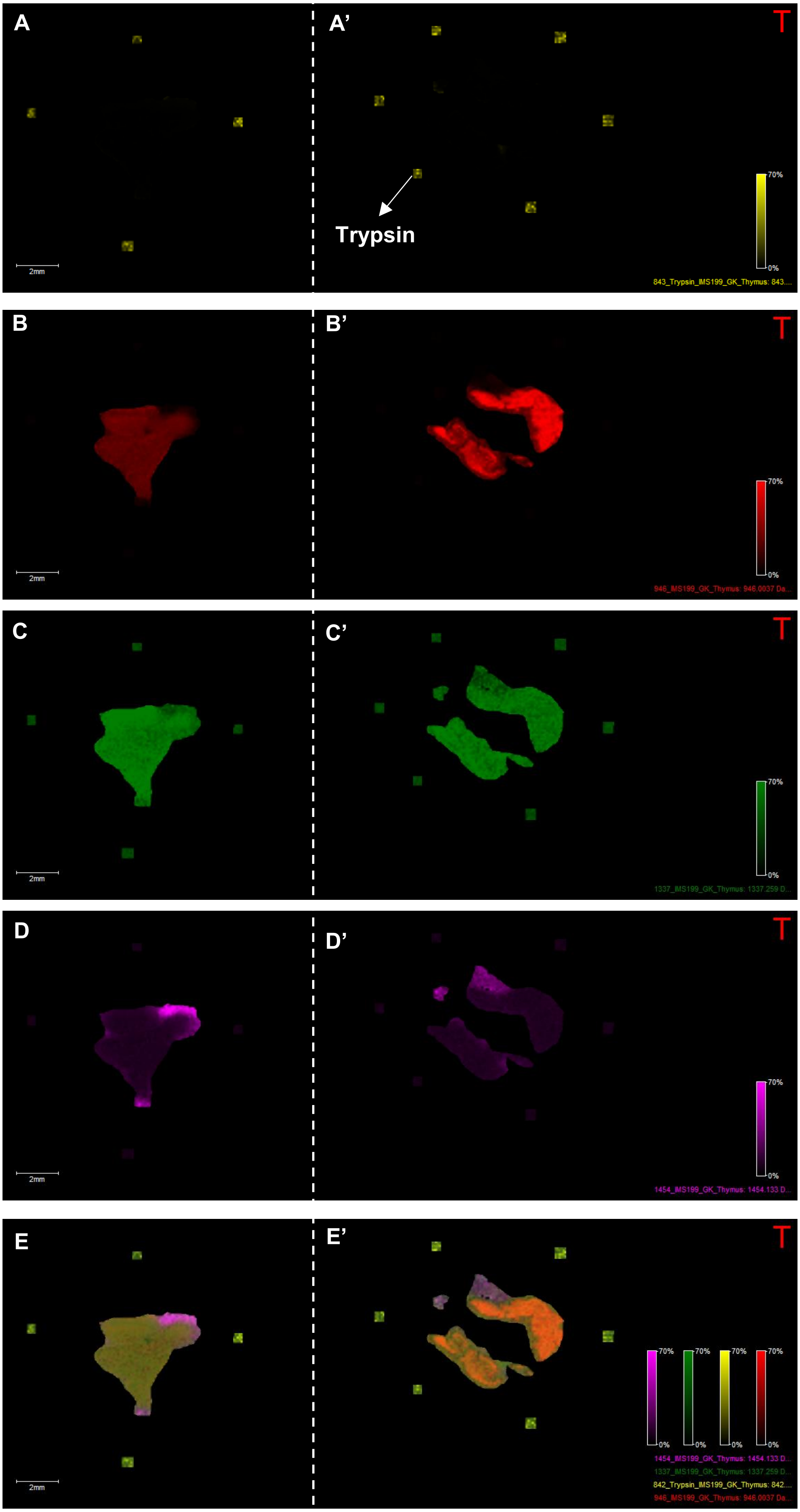


**Supplementary Figure S2. A and A':** The signal at 843 Da corresponds to trypsin (yellow). Three signals at **B and B':** 946 Da (red), **C and C':** 1337 Da (green) and **D and D':** 1454 Da (pink) belonging to three different clusters corresponding to three distinct thymic compartments. **E and E':** The overlay shows that the distinct locations of the three ion heat maps were retained.

### Figure S3

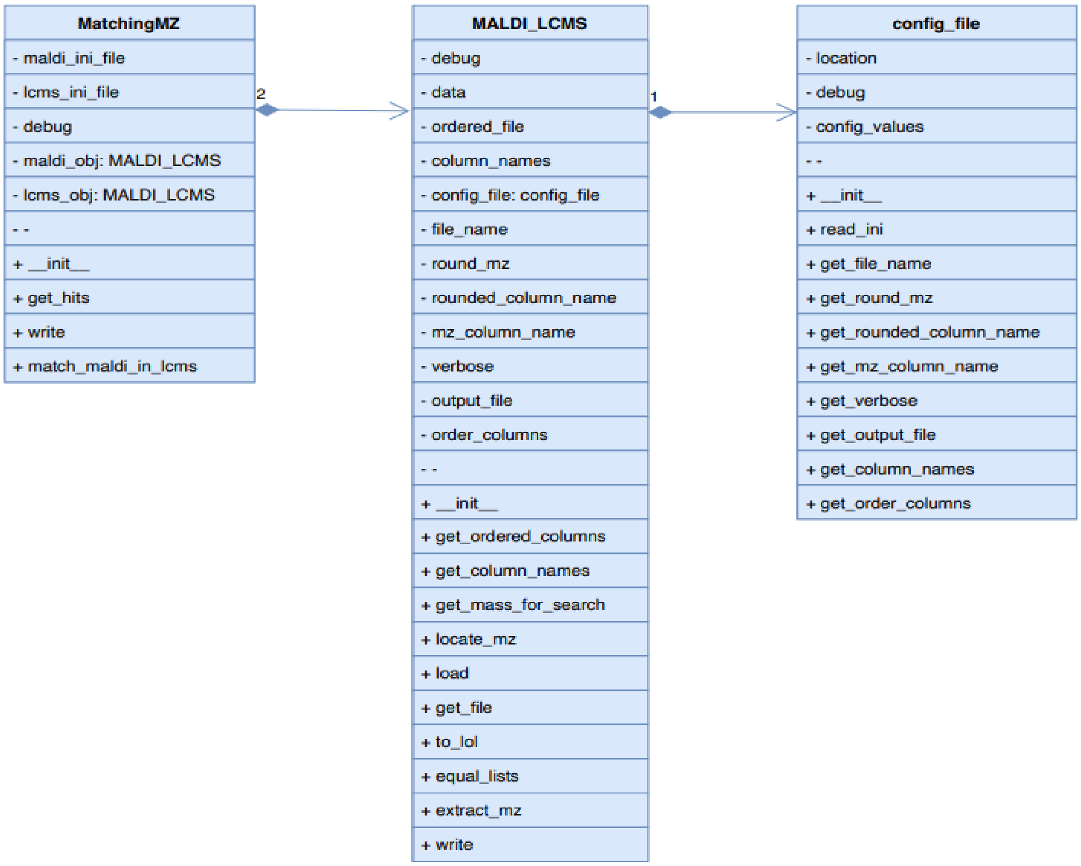


**Supplementary Figure S3.** This figure shows the pepBridge class diagrams. pepBridge is an in-house developed python based program (pepBridge.py) that can process .csv files that contain MALDI imaging m/z, MALDI intensities and the peptide table from the LC-MS/MS analysis for MALDI-MSI and LC-MS/MS peak matching.

### Figure S4

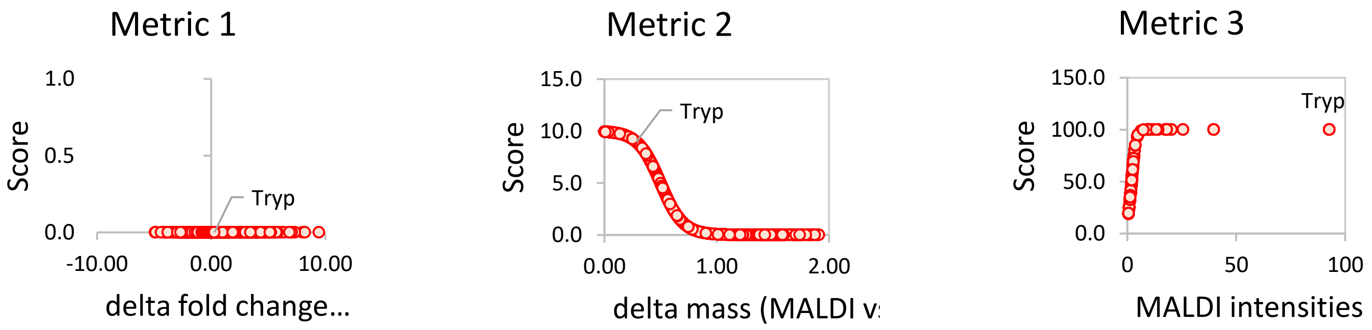


**Supplementary Figure S4.** Metrics for trypsin standard. Plot displaying the scores for all the signals matched in the spectra corresponding to the trypsin spotting. Data show how trypsin is highlighted as main hit (as expected) when MALDI and LC-MS data are combined.

### Figure S5

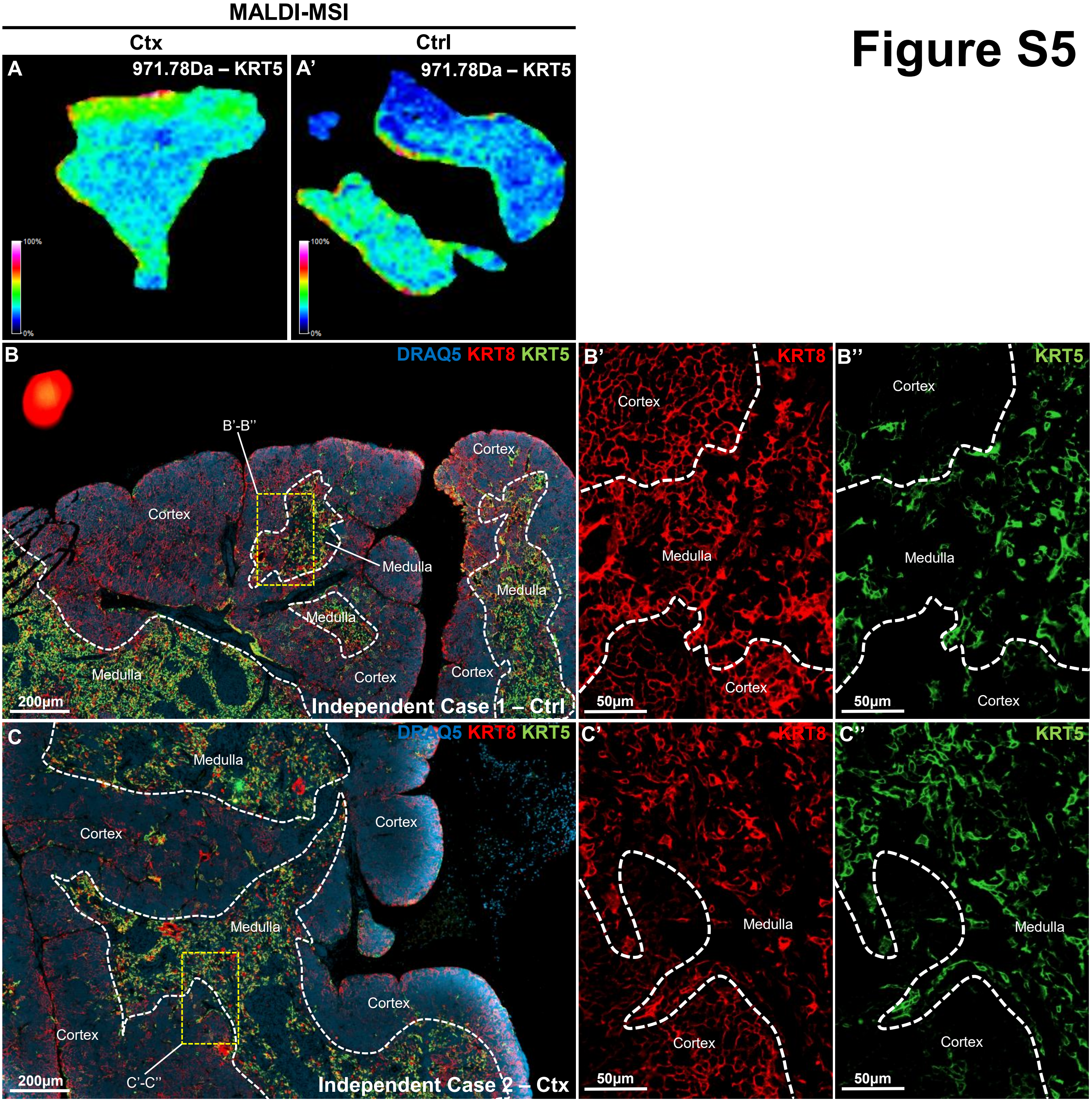


**Supplementary Figure S5.** MALDI vs immunohistochemistry comparison.

### Figure S6

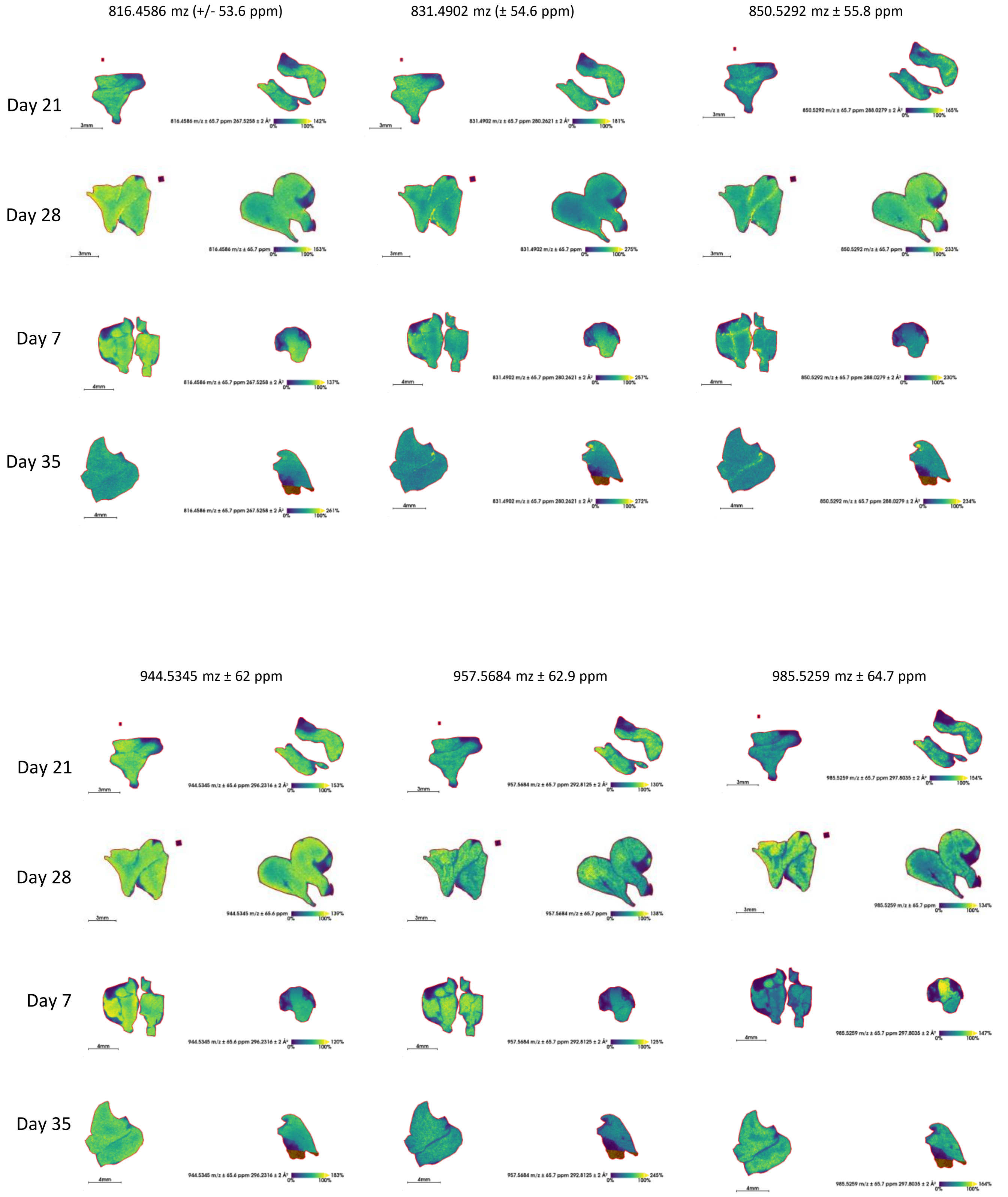


**Supplementary Figure S6.** Top 10 most abundant peptide ions (m/z) that showed the greatest difference between Ctrl and Ctx from imaging data acquired from the timsTOF flex. First part

### Figure S7

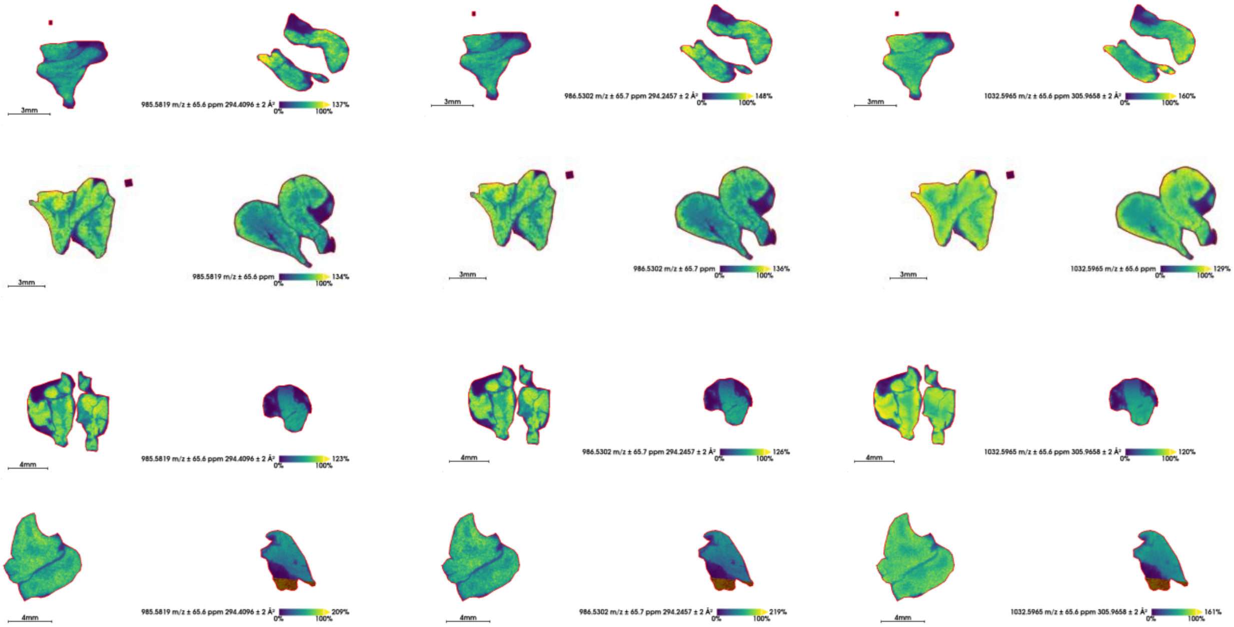


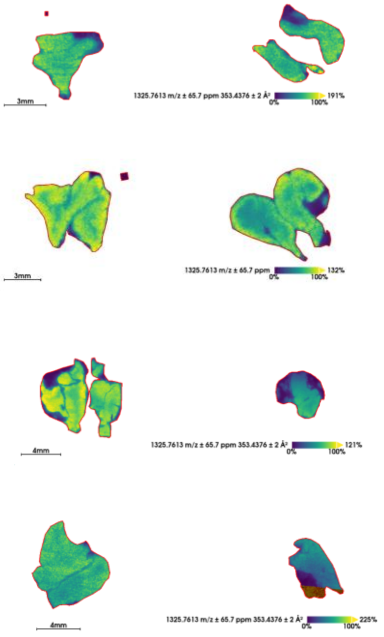


**Supplementary Figure S7.** Top 10 most abundant peptide ions (m/z) from timsTOF flex. Second part.
